# A low-cost recombinant glycoconjugate vaccine confers immunogenicity and protection against enterotoxigenic *Escherichia coli* infections in mice

**DOI:** 10.1101/2022.10.31.514630

**Authors:** Asher J. Williams, Katherine F. Warfel, Primit Desai, Jie Li, Jen-Jie Lee, Derek A. Wong, Sarah E. Sobol, Michael C. Jewett, Yung-Fu Chang, Matthew P. DeLisa

**Affiliations:** Robert F. Smith School of Chemical and Biomolecular Engineering, Cornell University, Ithaca, NY 14853 USA; Department of Chemical and Biological Engineering, Northwestern University, 2145 Sheridan Road, Technological Institute E136, Evanston, IL 60208, USA; Chemistry of Life Processes Institute, Northwestern University, 2170 Campus Drive, Evanston, IL 60208, USA; Center for Synthetic Biology, Northwestern University, 2145 Sheridan Road, Technological Institute E136, Evanston, IL 60208, USA; Biochemistry, Molecular and Cell Biology, Cornell University, Ithaca, NY 14853; Department of Population Medicine and Diagnostic Sciences, College of Veterinary Medicine, Cornell University, Ithaca, NY, 14853 USA; Cornell Institute of Biotechnology, Cornell University, Ithaca, NY 14853 USA

**Author notes:** Address correspondence to: Matthew P. DeLisa, Robert Frederick Smith School of Chemical and Biomolecular Engineering, Cornell University, Ithaca, NY 14853 USA.

## Abstract

Enterotoxigenic *Escherichia coli* (ETEC) is the primary etiologic agent of traveler’s diarrhea and a major cause of diarrheal disease and death worldwide, especially in infants and young children. Despite significant efforts over the past several decades, an affordable vaccine that significantly reduces mortality and morbidity associated with moderate to severe diarrhea among children under the age of 5 years remains an unmet aspirational goal. Here, we describe robust, cost-effective biosynthetic routes that leverage glycoengineered strains of non-pathogenic *Escherichia coli* or their cell-free extracts for producing conjugate vaccine candidates against two of the most prevalent O serogroups of ETEC, O148 and O78. Specifically, we demonstrate site-specific installation of O-antigen polysaccharides (O-PS) corresponding to these serogroups onto licensed carrier proteins using the oligosaccharyltransferase PglB from *Campylobacter jejuni*. The resulting conjugates stimulate strong O-PS-specific humoral responses in mice and elicit IgG antibodies that possess bactericidal activity against the cognate pathogens. We also show that one of the prototype conjugates decorated with serogroup O148 O-PS confers protection against ETEC infection in mice. We anticipate that our bacterial cell-based and cell-free platforms will enable creation of multivalent formulations with the potential for broad ETEC serogroup protection and increased access through low-cost biomanufacturing.

## INTRODUCTION

Enterotoxigenic *Escherichia coli* (ETEC) is one of the most common causes of diarrheal disease worldwide and the leading cause of traveler’s diarrhea, especially in locations where clean water and sanitation remain limited ^1, 2^. In addition to acute diarrhea-associated morbidity, ETEC is also one of the leading causes of mortality, disproportionately affecting children under the age of 5 years who lack immunity from prior exposure ^3, 4^. Deaths from infectious diarrhea have been on the decline for a few decades, due in part to the introduction of oral rehydration therapy. However, ETEC continues to wreak havoc in terms of acute morbidity and associated sequelae that compound the impact of infection in young children, including growth stunting, malnutrition, and impaired cognitive impairment ^5, 6^.

A vaccine that prevents both acute illness as well as the sequelae associated with ETEC infection has long been a priority of the World Health Organization (WHO) ^7^. At present, ETEC vaccine development efforts are primarily centered on inducing immune responses against a subset of important virulence factors including colonization factor/cell surface (CF/CS) antigens and enterotoxins, namely heat-stable toxin (ST) and heat-labile toxin (LT) ^8–11^. These factors comprise a classical model for ETEC molecular pathogenesis in which ETEC colonizes the small intestine using plasmid-encoded CF/CS antigens followed by production of one or more enterotoxin (*e.g*., LT, ST) that drive fluid export and diarrhea ^12^. Despite significant efforts over several decades, human trials involving vaccine candidates based on this classical paradigm have met limited success ^11, 13–15^. Hence, further improvements are needed to increase the levels of protection against more severe ETEC diarrhea and to expand protection to the breadth of ETEC serotypes. Consequently, there are still no licensed vaccines for ETEC.

To expand the repertoire of antigens that could be targeted in future vaccine designs, several studies have assessed adaptive immune responses in humans following either experimental challenge with ETEC or oral administration of an inactivated ETEC vaccine ^16–18^. A number of non-canonical antigens beyond the classic vaccine targets were identified, including secreted proteins (*e.g*., EatA, EtpA, and YghJ), cell surface-expressed proteins (*e.g*., Ag43, OmpW), and lipopolysaccharide (LPS), among others. Unlike the secreted and cell-surface protein antigens, LPS are glycolipids that include an outermost O-antigen polysaccharide (O-PS) component that is composed of repeating subunits that extend from the surface of the bacteria ^19^. Interestingly, the inactivated ETEC vaccine strain, ETVAX, elicited response frequencies against serogroup O78 LPS (which is expressed on the strain) that were comparable to or higher than those against the vaccine CFs in infants. This suggests that LPS is a potent antigen that may contribute to vaccine-induced protection ^17^. Other studies have indicated that O-PS is common among ETEC strains associated with diarrheal illness ^20, 21^, with more than 78 O serogroups identified in ~1,000 ETEC isolates from widespread locations ^20^. While this number is impractically high for developing a broadly protective, multivalent vaccine, 10 of these serogroups (O6, O8, O9, O25, O27, O78, O128, O148, O153 and O159) account for >75% of the isolates, suggesting that a 10-valent polysaccharide vaccine could afford broad protection with the fewest components possible. At present, however, virtually no attention has been paid to ETEC LPS/O-PS as a subunit vaccine antigen.

One barrier to the development of an LPS/O-PS-based vaccines in general is the fact that purified polysaccharides, while partially immunogenic in adults, are often completely incapable of eliciting an antibody response in infants and children, the population for whom an ETEC vaccine is most needed. This problem results from an inability of polysaccharides to interact with the receptors on T cells, but can be solved by covalently coupling the LPS or O-PS structure to a CD4^+^ T cell-dependent antigen such as an immunogenic protein carrier ^22^. The resulting conjugates invoke a T-cell response that results in strong polysaccharide-specific antibody responses, immunological memory, and high immunogenicity in young children ^23^. Indeed, conjugates are a highly efficacious and safe strategy for protecting against virulent pathogens, with successful vaccines licensed worldwide against *Haemophilus influenzae*, *Neisseria meningitidis* serogroups (tetravalent), *Streptococcus pneumoniae* (up to 20-valent), and *Salmonella typhi*, and many others in clinical development ^23^.

Despite their effectiveness, traditional conjugate vaccines are not without their drawbacks. Most notable among them is the complex, multistep process required for the purification, isolation, and conjugation of bacterial polysaccharides, which is expensive, time consuming, and low yielding ^24^. A greatly simplified and cost-effective alternative involves metabolic engineering of bacteria to generate recombinant strains that serve as mini factories for one-step production of an unlimited and renewable supply of pure conjugates ^25^. This bioconjugation approach is based on engineered protein glycosylation in non-pathogenic *E. coli* strains, wherein an O-PS molecule is conjugated to a co-expressed carrier protein by the *Campylobacter jejuni* oligosaccharyltransferase (OST) PglB (*Cj*PglB) ^26^. To date, several unique conjugates have been produced by this method, with a few currently under clinical investigation ^25^. Building on these cell-based efforts, we recently described a method called iVAX (*in vitro* conjugate vaccine expression) that enables conjugate vaccine biosynthesis using cell-free extracts derived from glycosylation-competent *E. coli* strains ^27^. The iVAX platform promises to accelerate vaccine development and enable decentralized, cold chain–independent biomanufacturing by using cell lysates, rather than living cells, to make conjugates *in vitro*.

In the current study, we describe two biosynthetic routes – one cell-based and the other cell-free – for low-cost production of conjugate vaccine candidates against two of the most prevalent O serogroups of ETEC, O148 and O78 ^20^. Both routes enabled site-specific installation of ETEC O-PS onto carrier proteins used in licensed vaccines, namely the nonacylated form of protein D (PD) from *Haemophilus influenzae* and cross-reactive material 197 (CRM_197_), a genetically detoxified variant of the *Corynebacterium diphtheriae* toxin (DT). The resulting conjugates stimulated strong O-PS-specific IgG antibody titers in mice, with the resulting antibodies possessing bactericidal activity against the cognate pathogens. For one of the prototype conjugates decorated with serogroup O148 O-PS, we further demonstrated protective efficacy against ETEC infection in mice. Overall, our work expands the inventory of antigens for ETEC vaccine design and provides an important first step towards the creation of a custom, multivalent vaccine with potential for broad ETEC coverage and increased access through adoption of simplified, low-cost biomanufacturing platforms.

## RESULTS

### Expression of ETEC serogroup O148 O-PS antigen in non-pathogenic *E. coli* cells

Biosynthesis of the O-PS antigen from ETEC serogroup O148 involved plasmid pMW07-O148 ^28, 29^, which encodes the 10.2 kb O-PS gene cluster from ETEC strain B7A (serotype O148:H28) ^30^ (**Fig. 1a**). To confirm O-PS expression, we took advantage of the fact that O-PS antigens assembled in the cytoplasmic membrane of *E. coli* cells are transferred onto lipid A-core by the O-antigen ligase, WaaL. The lipid A-core-linked O-PS molecules are then shuttled to the outer membrane, becoming displayed on the cell surface where they are readily detectable with antibodies or lectins having specificity for the O-PS structure. As expected, non-pathogenic *E. coli* W3110 cells, which carry a copy of the *waaL* gene, were observed to express the ETEC O-PS antigen on their surface as evidenced by cross reactivity of nitrocellulose-spotted cells with an anti-ETEC O148 antibody (**Supplementary Fig. 1a**). The binding observed for these cells was on par with that measured for ETEC strain B7A ^31^, which natively expresses the O148 O-PS antigen. In contrast, both plasmid-free W3110 cells and W3110 cells carrying empty pMW07 plasmid showed little to no cross reactivity. A similar lack of cross reactivity was observed for CLM24 cells, which have a deletion in the *waaL* gene, confirming that the recombinant O-PS antigen was assembled via the canonical lipid A-core pathway.

**Figure 1.**
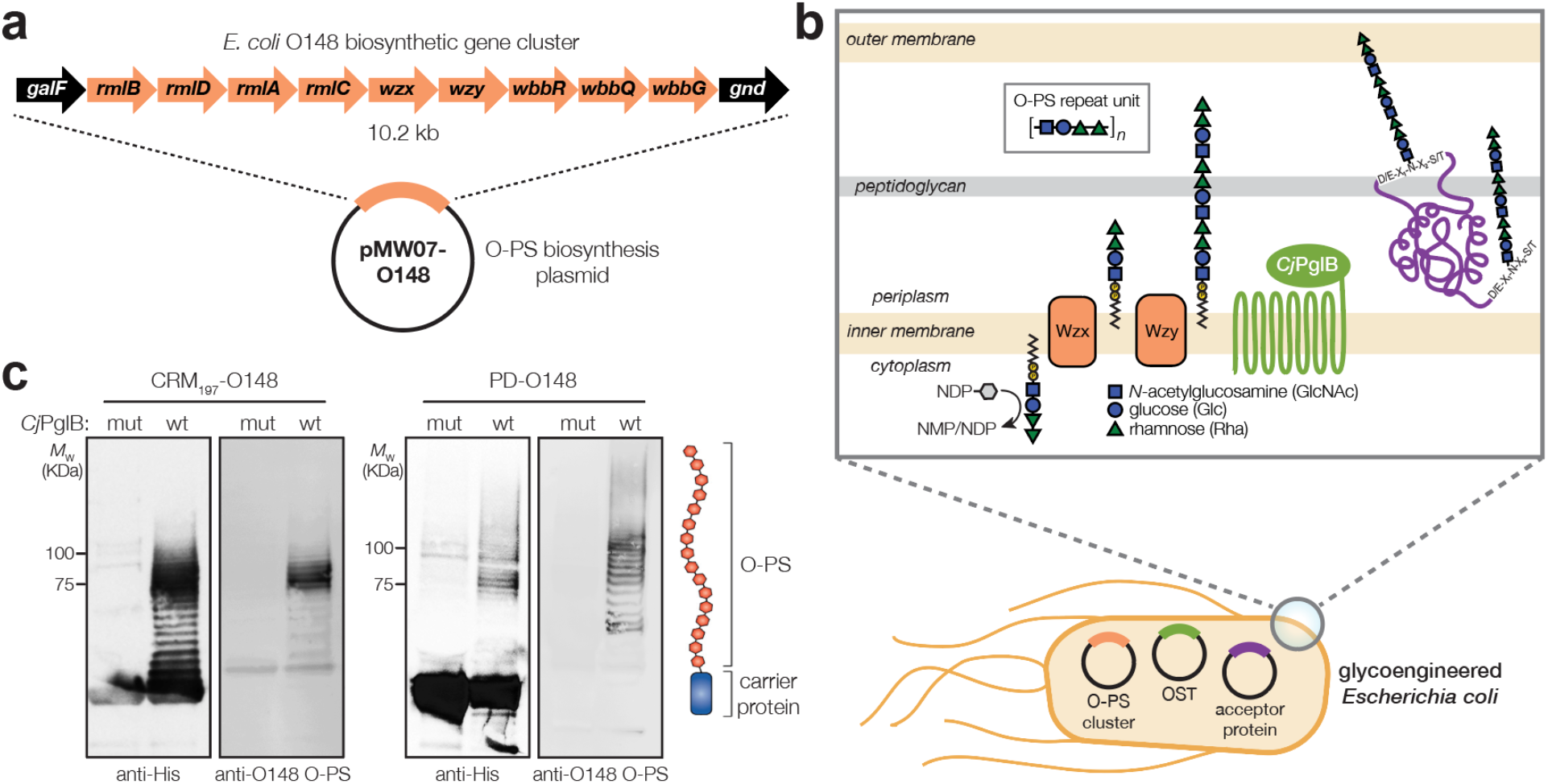
Biosynthesis of glycoconjugate vaccine candidates against ETEC bacteria. (a) Biosynthesis of ETEC O148 O-PS from plasmid pMW07-O148, which encodes the entire O-PS locus from ETEC strain B7A (serotype O148:H28) between *galF* and *gnd*. (b) Glycoconjugate biosynthesis is enabled by combining engineered O-antigen biosynthesis with *N*-linked protein glycosylation by *C. jejuni* PglB (*Cj*PglB). Several plasmid-encoded glycosyltransferases sequentially add the repeat-unit sugars to the lipid carrier undecaprenol pyrophosphate in the cytoplasmic membrane. The lipid-linked oligosaccharide is flipped by Wzx and polymerized by the Wzy polymerase, and subsequently transferred to ‘DQNAT’ acceptor sites in the carrier protein by PglB. Deletion of *waaL* in the host strain is used to eliminate transfer of the O-PS to lipid A-core and ensure efficient transfer to the acceptor protein, while deletion of *lpxM* results in pentaacylated lipid A structure having significantly reduced toxicity. (c) Immunoblot analysis of purified carrier proteins derived from *E. coli* CLM24 cells carrying a plasmid encoding either CRM_197_^4xDQNAT^ or PD^4xDQNAT^ along with plasmid pMW07-O148 encoding the ETEC O148 O-PS biosynthetic pathway and plasmid pMAF10 encoding wild-type *Cj*PglB (wt) or an inactive mutant of *Cj*PglB (mut) as indicated. Blots were probed with anti-His antibody to detect acceptor proteins and anti-ETEC O148 antibody to detect O-PS. Images depict aglycosylated and multiply glycosylated forms of CRM_197_^4xDQNAT^ or PD^4xDQNAT^. Molecular weight (*M*_W_) markers are indicated on the left. Results are representative of three biological replicates.

### Glycosylation of licensed vaccine carrier proteins with ETEC O148 O-PS

To generate a strong IgG response and lasting immunity, it is desirable to employ a highly immunogenic protein as a carrier for the polysaccharide antigen, in this case ETEC serogroup O148 O-PS. To this end, we sought to engineer non-pathogenic *E. coli* with the ability to glycosylate a set of carrier proteins, namely PD and CRM_197_, that are currently used in licensed conjugate vaccines (**Fig. 1b**). To enable conjugation of O-PS antigens to these carrier proteins, both were modified at their C termini with four tandem repeats of an optimized bacterial glycosylation motif, DQNAT ^32^, followed by a 6x-His tag to enable detection via Western blot analysis and purification by Ni-NTA chromatography. A signal peptide sequence derived from the *E. coli* DsbA protein was fused to the N-terminus to localize CRM_197_ and PD to the periplasm in a manner that is compatible with *N*-linked glycosylation ^33^. Each of the resulting plasmids, pTrc99A-CRM_197_^4xDQNAT^ and pTrc99A-PD^4xDQNAT^, were used to transform *E. coli* strain CLM24 carrying plasmid pMW07-O148 that encoded the O-PS biosynthetic enzymes and plasmid pMAF10 that encoded *Cj*PglB ^26^. CLM24 cells were used because they have a deletion of the gene encoding the WaaL O-antigen ligase that makes undecaprenol pyrophosphate (UndPP)-linked glycans including O-PS structures exclusively available to *Cj*PglB by preventing their unwanted transfer to lipid A-core ^26^.

Following overnight expression of *Cj*PglB along with either CRM_197_^4xDQNAT^ or PD^4xDQNAT^ in the presence of the ETEC serogroup O148 biosynthetic enzymes, cells were lysed, and His-tagged carrier proteins were purified by Ni-NTA chromatography. Elution fractions from each sample were separated by SDS-PAGE and subjected to immunoblotting using an anti-His antibody to detect the carrier proteins or antiserum specific for ETEC O148 LPS to detect the O-PS antigen. This analysis revealed that both CRM_197_^4xDQNAT^ and PD^4xDQNAT^ were readily glycosylated with ETEC O148 O-PS glycans (**Fig. 1c**). Importantly, we observed a ladder-like banding pattern for both O148 O-PS-linked carrier proteins (hereafter CRM_197_-O148 and PD-O148), which is characteristic of *Cj*PglB-mediated O-PS transfer ^26^ and results from variability in the chain length of O-PS antigens generated by the Wzy polymerase ^19^. The most intense laddering signal was observed above 75 kDa, suggesting that the carriers were heavily decorated with ETEC O148 O-PS structures comprised of >10 repeating units (RUs). Control reactions with CLM24 cells that lacked the O-PS biosynthetic plasmid or expressed a catalytically inactive *Cj*PglB enzyme, generated by introducing D54N and E316Q substitutions ^34^, confirmed that O-PS conjugation to the carrier proteins depended on both the O-PS biosynthetic enzymes and *Cj*PglB (**Fig. 1c**; shown for mutant *Cj*PglB).

### Immunogenicity of PD-O148 glycoconjugate in mice

To investigate conjugate immunogenicity, BALB/c mice were immunized subcutaneously (s.c.) with glycosylated PD bearing the ETEC O148 O-PS antigen and serum from these animals was analyzed by enzyme-linked immunosorbent assay (ELISA) to determine antibody titers. BALB/c mice were immunized with 50 μg doses of protein, either PD alone or PD-O148 conjugate, adjuvanted with aluminium phosphate, and subsequently injected with identical booster doses at 21 and 42 days after the initial injection (**Fig. 2a**). Upon analyzing sera collected on day 56, we found that BALB/c mice receiving the PD-O148 conjugate produced high titers of IgG antibodies that specifically recognized LPS derived from ETEC strain B7A (**Fig. 2b**). These serum IgG levels were significantly elevated (~2 orders of magnitude) compared to the titers measured in sera of control mice receiving buffer or aglycosylated PD. The ability of the PD-O148 conjugate to elicit strong IgG titers against ETEC B7A LPS provides further validation for using non-pathogenic *E. coli* strains engineered with protein glycosylation machinery as hosts for conjugate vaccine production.

**Figure 2.**
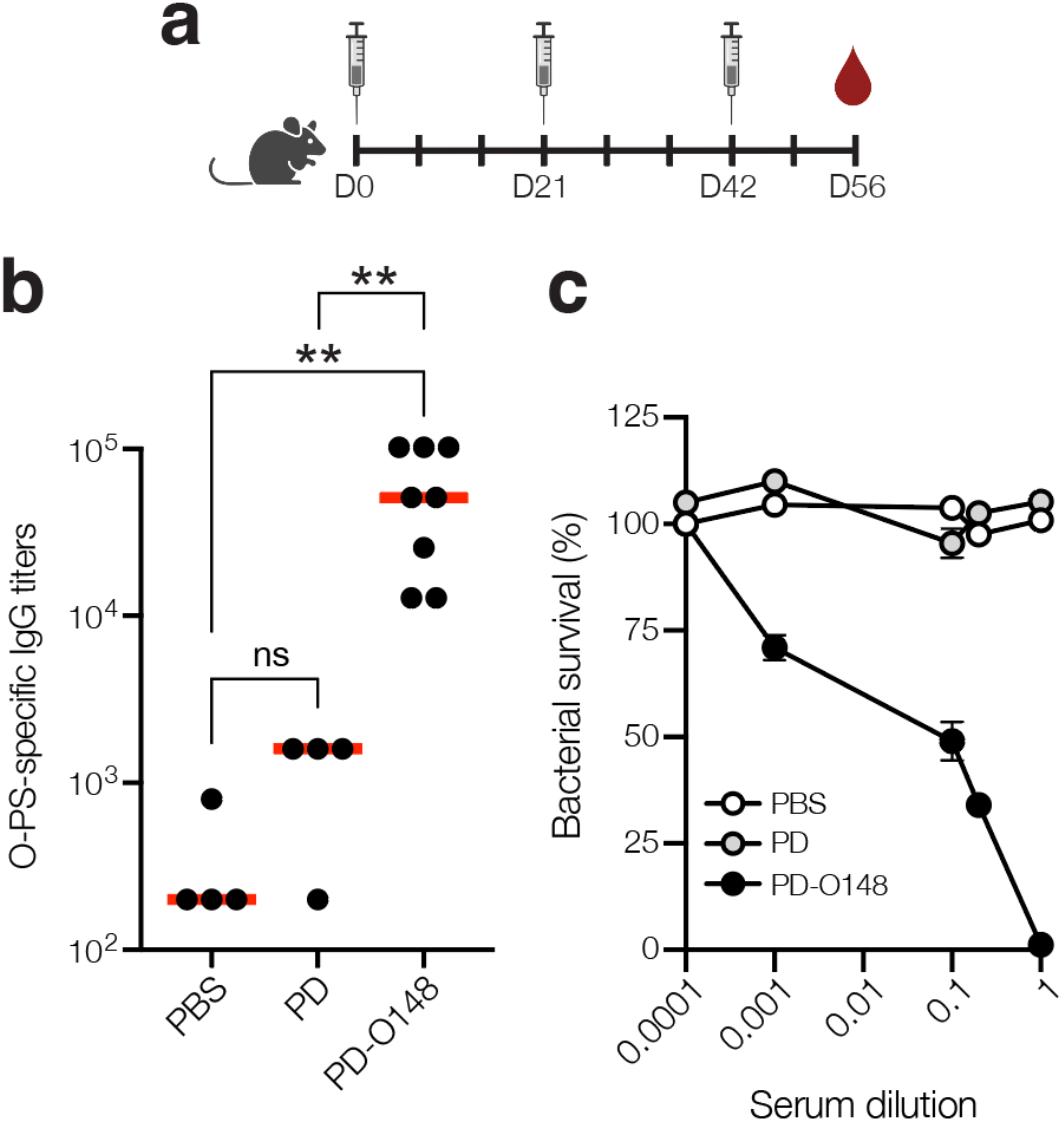
Glycoconjugates are immunogenic. (a) Schematic of the prime-boost immunization schedule. Mice received an initial injection on day 0 and identically formulated booster injections on days 21 and 42. Blood was drawn on days 0, 35, 49, and at study termination on day 56. (b) O-PS-specific IgG titers in day 56 serum of individual mice (black circles) and mean titers of each group (red lines) as determined by ELISA with LPS derived from ETEC strain B7A (serotype O148:H28) as immobilized antigen. Groups of three BALB/c mice were immunized s.c. with 100 μL PBS alone or PBS containing 50 μg of either aglycosylated PD carrier protein or glycosylated ETEC O148 conjugate (PD-O148) adjuvanted with aluminium phosphate adjuvant. Mice were boosted on days 21 and 42 with identical doses of each immunogen. Statistical significance was determined by unpaired *t* test with Welch’s correction (\**p* < 0.05; \*\**p* < 0.01; ns, not significant). (c) Bactericidal killing activity of antibodies in the serum of mice immunized with PBS, unmodified PD carrier protein, or ETEC O148 conjugate (PD-O148). Survival data were derived from a standard SBA where dilutions of serum from immunized mice were tested against ETEC strain B7A (serotype O148:H28) in the presence of human complement. SBA curves were generated with pooled sera from each group. Data are the mean of three biological replicates ± SEM. efficacy and protectiveness of our glycoengineered vaccine candidate.

### Functionality of ETEC O-PS-specific serum antibodies

Next, PD-O148 conjugate vaccine-induced serum antibodies were evaluated for the ability to promote complement-mediated killing of ETEC strain B7A by serum bactericidal assay (SBA). SBA is an established method by which the activity of IgGs against bacterial pathogens can be measured. It often correlates with protection for serotypes of a pathogen and is a key *in vitro* method for measuring the functional activity of antibodies ^35^. Of relevance here, several groups have developed bactericidal assays for evaluating whether serum IgG antibodies can potentiate the killing of different ETEC strains ^36, 37^. Using a similar methodology, we investigated the idea that O-PS-specific serum IgGs elicited by the PD-O148 glycoconjugate would bind to the corresponding LPS on the surface of ETEC strain B7A and, in the presence of components of the human complement system, would mediate bacteriolysis of the enteric pathogen. For the sera derived from mice vaccinated with the PD-O148 conjugate, ~50% survival of ETEC B7A cells (corresponding to ~50% killing activity) was observed at dilutions as high as 10-fold (**Fig. 2c**). In contrast, virtually no killing was observed for sera derived from mice treated with PBS or the aglycosylated PD carrier protein as evidenced by pathogen survival that was close to 100%. Additionally, near complete killing was observed for the undiluted sera of vaccinated mice, whereas no killing was observed for the PBS and aglycosylated PD groups at the same serum dilution. These results confirm the immunological functionality of PD-O148 conjugate vaccine-induced IgGs present in the sera of immunized mice and predict the

### Protective efficacy of PD-O148 glycoconjugate in mice

Encouraged by the SBA results, we tested the ability of the PD-O148 conjugate to protect mice in a murine model of an orally administered ETEC B7A infection. A major hurdle in developing enteric vaccines is the lack of a suitable small animal model to study the efficacy and immunogenicity of potential ETEC vaccines prior to testing in larger animals or humans. Nevertheless, ETEC infection is commonly induced by the pathogen via oral administration, with mice becoming colonized with ETEC following oral challenge using inocula as small as 10^3^ colony-forming units (CFUs) ^38^. Oral gavage was selected as an infection model as it closely reflects the route of infection and potentially recapitulates relevant outcomes of ETEC infection seen in humans ^39^. In this study, mice were pretreated with the antibiotic streptomycin, allowing them to more closely mimic the disease symptoms that are often seen in humans ^39, 40^. Following immunization according to an identical schedule as above, mice were infected by oral gavage with ~1×10^4^ CFUs of ETEC strain B7A (**Fig. 3a**). After challenge infection, mice were monitored for signs and symptoms of enteric illness (*e.g*., water diarrhea). Diarrhea induced by ETEC infection was greatly reduced in the PD-O148-vaccinated group overall three days post-infection, which was in stark contrast to buffer and aglycosylated PD control groups that developed watery diarrhea. We also collected stool samples to detect shedding of the challenge strain. Diarrheal illness, when it occurs, is associated with higher fecal shedding levels of ETEC in animals and humans ^39, 41^. Post-challenge shedding levels of the challenge strain ETEC B7A was detected using quantitative PCR (qPCR) to examine fecal pellet DNA extracts for the presence of LT encoded by the *eltA* gene. Consistent with the observed protection against diarrhea, only mice receiving the PD-O148 conjugate exhibited a significant reduction in ETEC stool shedding detected at day 3 post-infection (~2-log reduction compared to the buffer and aglycosylated PD carrier protein control groups) (**Fig. 3b**). Taken together, these results clearly demonstrated the ability of the PD-O148 conjugate vaccine candidate to confer protection against ETEC infection in mice, consistent with the conjugate’s ability to stimulate high titers of IgG antibodies that possess potent bactericidal activity.

**Figure 3.**
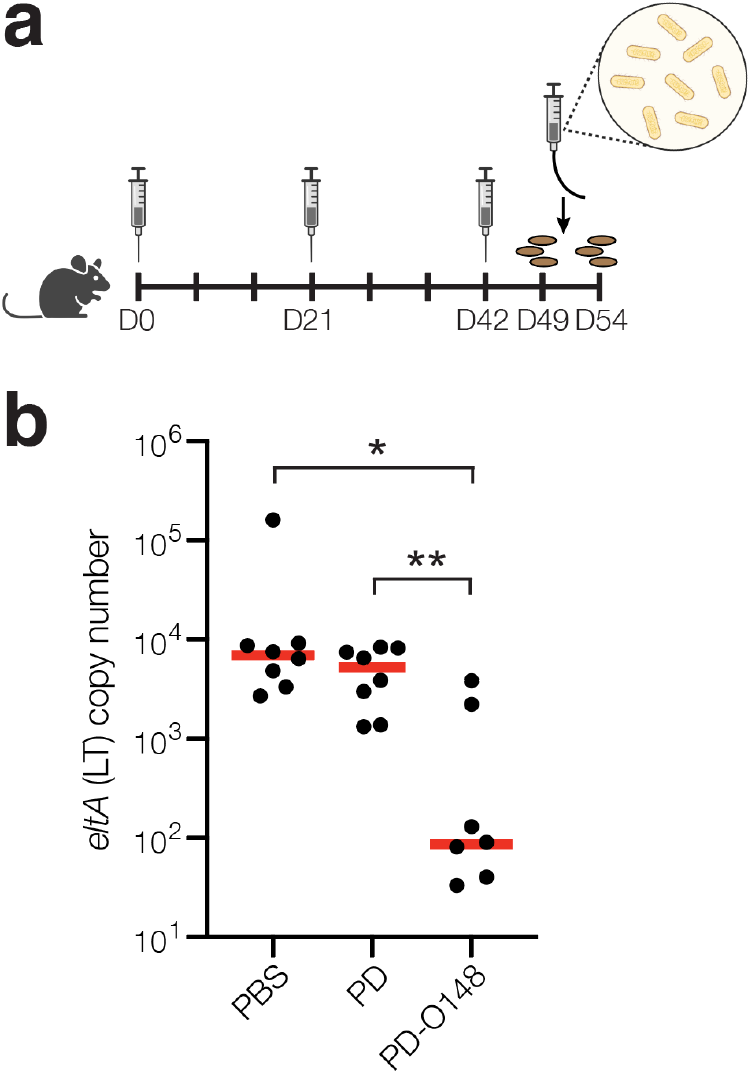
Glycoconjugates are functional and protective. (a) Schematic of the immunization and challenge schedule. Mice received an initial immunogen injection on day 0 and identically formulated booster injections on days 21 and 42. On day 49, mice received antibiotics in the drinking water to eradicate normal flora and fecal pellets were collected. On day 51, mice were subsequently infected by gavage and checked daily for 3 days. On day 54, fecal pellets were collected, and mice were sacrificed. (b) Vaccine effects on stool shedding of ETEC B7A in infected mice 3 days post-infection. Groups of 8 BALB/c mice receiving PBS, unmodified PD carrier protein, or PD-O148 were infected with ~1 × 10^4^ CFUs of ETEC B7A. Fecal samples were collected for DNA extraction and analyzed by qPCR to detect pathogen. ETEC burden in fecal pellets was measured by qPCR of the *eltA* gene, which encodes heat-labile endotoxin (LT). Data are the mean (red bars) of individual mice (black circles). Statistical significance was determined by unpaired *t* test with Welch’s correction (\**p* < 0.05; \*\**p* < 0.01).

### Biosynthesis and immunogenicity of an ETEC serogroup O78-directed conjugate

As a first step towards such a multivalent formulation, we sought to create a glycoconjugate vaccine candidate against ETEC serogroup O78 by implementing an identical strategy as outlined above for serogroup O148. We chose serogroup O78 because it is one of the most widely distributed and most frequently occurring ETEC serogroups that, together with ETEC serogroups O6, O8, O27, O148, O153 and O169, accounts for nearly 25% of the nearly 1,000 isolated identified worldwide ^20^. Introduction of plasmid pMW07-O78 ^27, 29, 42^ (**Supplementary Fig. 2a**) into non-pathogenic *E. coli* strain W3110 enabled expression of cell-surface O78 O-PS molecules that were readily detected by dot blot analysis (**Supplementary Fig. 1b**). By combining plasmid pMW07-O78 together with the plasmids for expressing *Cj*PglB and the CRM_197_ carrier protein in CLM24 Δ*lpxM* cells, we were able to produce glycosylated CRM_197_ bearing the O78 O-PS antigen (**Supplementary Fig. 2b**). The glycosylated CRM_197_-O78 conjugate was used to immunize BALB/c mice and was observed to induce high titers of IgG antibodies that specifically recognized LPS derived from ETEC strain H10407 (serotype O78:H11) ^43^ (**Supplementary Fig. 2c**). Collectively, these results highlight the modularity of the biosynthetic approach, enabling facile production of an additional serogroup-specific conjugate simply by swapping out the O-PS biosynthesis plasmid. Such interchangeability will be key to generating a multivalent conjugate formulation that confers broad ETEC serogroup protection, with the results here representing an important first step in that direction.

### Cell-free biosynthesis of ETEC O78 O-PS conjugate

In parallel to using living cells, we also explored whether a cell-free protein synthesis (CFPS) approach could be used to produce glycoengineered conjugate vaccine candidates against ETEC. To this end, we leveraged a modular technology for *in vitro* conjugate vaccine expression (iVAX) in portable, freeze-dried lysates from detoxified, non-pathogenic *E. coli* ^27^ (**Fig. 4a**). Previous studies demonstrated that iVAX reactions are capable of synthesizing clinically relevant doses of protective conjugate vaccines comprised of pathogen-specific O-PS antigens linked to licensed carrier proteins. Here, we produced an iVAX lysate from *E. coli* CLM24 Δ*lpxM* cells expressing the ETEC O78 O-PS biosynthetic pathway and *Cj*PglB. This lysate, which contained lipid-linked ETEC O78 O-PS and active *Cj*PglB, was used to catalyze iVAX reactions primed with plasmid DNA encoding the PD^4xDQNAT^ carrier protein. The products of these reactions were immunoblotted with anti-His antibody or a commercial anti-ETEC O78 antibody specific to the ETEC O78 O-PS. Similar to cell-based expression, cell-free iVAX reactions produced PD^4xDQNAT^ that was clearly glycosylated with the O78 O-PS antigen and exhibited the characteristic ladderlike banding pattern associated with O-PS chain-length variability (**Fig. 4b**). Control reactions with lysates from cells lacking the ETEC O78 O-PS were devoid of any detectable glycosylation. Following immunization of BALB/c mice with the iVAX-derived conjugate, we observed strong induction of IgG antibodies that specifically recognized LPS derived from ETEC strain H10407 (serotype O78:H11) (**Fig. 4c**), with serum IgG titers comparing favorably to those observed following immunization with the CRM_197_-O78 conjugate that was made in living cells. Finally, PD-O78 conjugate vaccine-induced serum antibodies were evaluated for the ability to promote complement-mediated killing of ETEC strain H10407 by SBA. Greater than ~50% killing activity of ETEC H10407 cells was observed for the sera derived from PD-O78-vaccinated-mice at dilutions as high as 10-fold, whereas no measurable killing was observed for sera derived from mice treated with PBS or aglycosylated PD (**Fig. 4d**).

**Figure 4.**
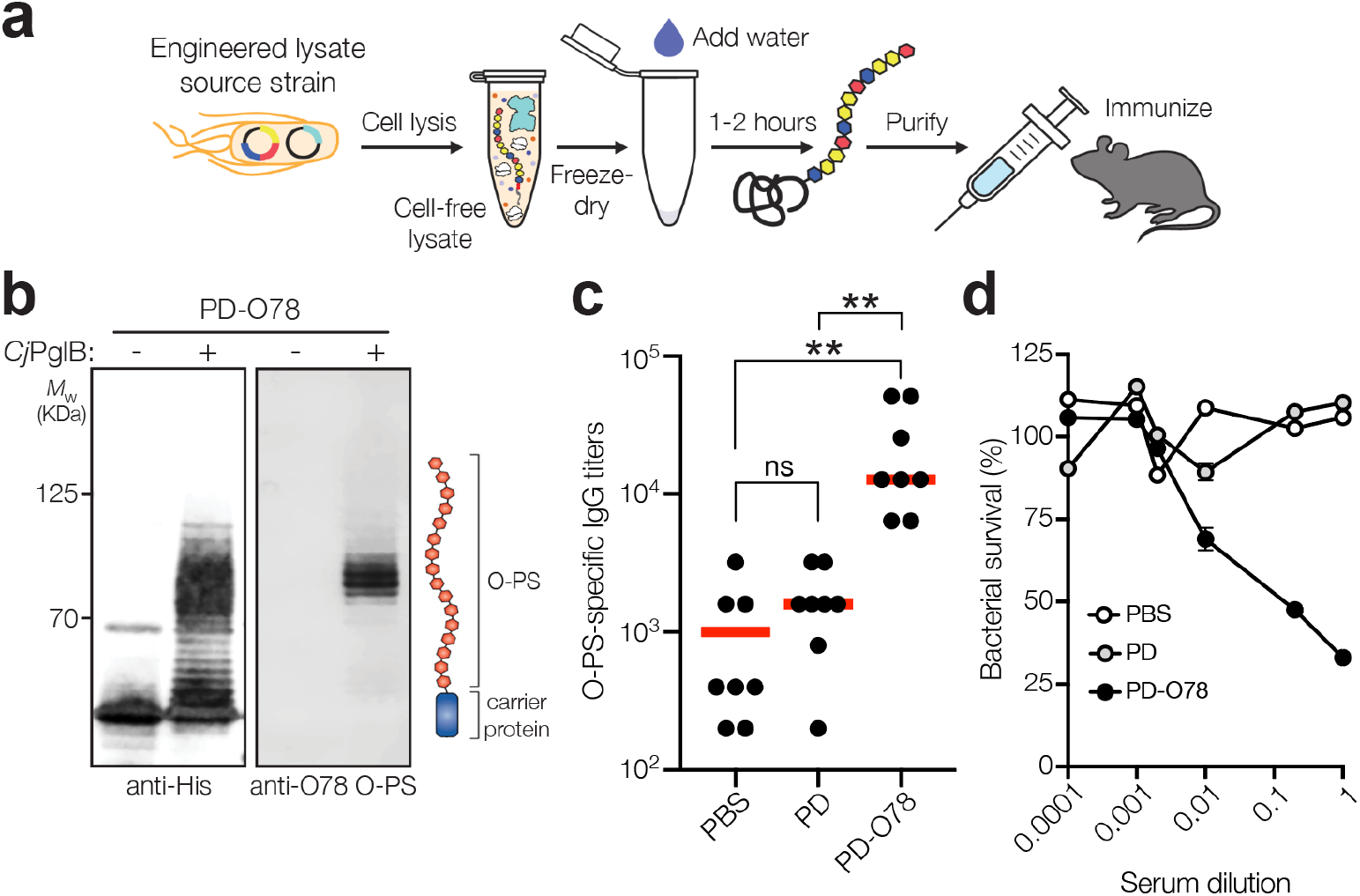
Glycoconjugates are functional and protective. (a) The iVAX platform provides a rapid means to develop and distribute conjugate vaccines against bacterial pathogens. (b) Immunoblot analysis of PD^4xDQNAT^ generated using iVAX lysate. Specifically, 5-mL reactions with glyco-enriched S12 extract derived from CLM24 Δ*lpxM* cells carrying pMW07-O78 and pSF-*Cj*PglB-LpxE were primed with plasmid pJL1-PD^4xDQNAT^. Blots were probed with anti-His antibody to detect acceptor proteins and anti-ETEC O78 antibody to detect O-PS antigens. Images depict aglycosylated and multiply glycosylated forms of PD^4xDQNAT^. Molecular weight (*M*_W_) markers are indicated on the left. Results are representative of three biological replicates. (c) O-PS-specific IgG titers in day 56 serum of individual mice (black circles) and mean titers of each group (red lines) as determined by ELISA with LPS derived from ETEC strain H10407 (serotype O78:H11) as immobilized antigen. Groups of three BALB/c mice were immunized s.c. with 100 μL PBS alone or PBS containing 24 μg of either aglycosylated PD carrier protein or glycosylated ETEC O148 conjugate adjuvanted with aluminium phosphate adjuvant. Mice were boosted on days 21 and 42 with identical doses of each immunogen. Statistical significance was determined by unpaired *t* test with Welch’s correction (\**p* < 0.05; \*\**p* < 0.01; ns, not significant). (d) Bactericidal killing activity of serum antibodies from mice immunized with same immunogens as in (c). Survival data were derived from a standard SBA where serum dilutions were tested against ETEC strain H10407 in the presence of human complement. SBA curves were generated with pooled sera from each group. Data are the mean of three biological replicates ± SEM.

## DISCUSSION

In the present study, we describe robust cell-based and cell-free bioconjugation strategies for producing conjugate vaccine candidates against two widespread O serogroups of ETEC, O148 and O78 ^20^. These strategies leveraged glycoengineered strains of non-pathogenic *E. coli* and their cell-free extracts for site-specific installation of two different ETEC O-PS structures onto the PD and CRM_197_ carrier proteins that are used in licensed vaccines. The resulting PD- and CRM_197_-based conjugates were strongly immunogenic in mice, eliciting high titers of O-PS-specific IgG antibodies that also exhibited potent bactericidal activity against ETEC strains B7A (serotype O148:H28) and H10407 (serotype O78:H11). In proof-of-concept challenge studies, mice vaccinated with the PD-O148 conjugate were protected against ETEC infection as evidenced by the lack of overt water diarrhea and significantly decreased ETEC stool shedding. Overall, our findings expand the repertoire of available ETEC antigens to include molecules that have largely been overlooked for subunit vaccine engineering. Moreover, the cell-based and cell-free platforms described here lay the foundation for future creation of a custom, multivalent vaccine with the potential for broad ETEC coverage and increased access through adoption of simplified, low-cost biomanufacturing platforms.

The immunogenicity and protective efficacy of our conjugates substantiate the use of ETEC O-PS as a subunit vaccine antigen. This antigenic expansion is significant for several reasons. First, even though glycoconjugate vaccines are one of the safest and most effective methods for preventing bacterial infections, there are surprisingly few that have been fully licensed. That said, these numbers are poised to increase in the coming years as cell-based and cell-free technologies for glycoengineering recombinant vaccines, such as those described here and elsewhere, ^25^ reach full maturity. As part of this maturation, it is imperative to continue growing the pipeline with as many promising anti-bacterial vaccine candidates as possible, especially as diarrheal and other vaccine-preventable diseases continue to be unmet challenges and as drug-resistant bacteria are predicted to threaten up to 10 million lives per year by 2050 ^44^. Second, all ETEC vaccines to enter clinical testing thus far are focused on the classical paradigm of ETEC pathogenesis and attempt to induce immune responses to select colonization factor (CF/CS) antigens and LT ^8–11^. However, the risk of focusing on the classical paradigm is that it constrains ETEC vaccinology to a subset of canonical antigens, which could be problematic if our current view of ETEC pathogenesis is incomplete, as has been suggested recently ^10^. For this reason, we decided to investigate O-PS from ETEC serogroups O148 and O78 with the goal of validating these non-canonical targets and supplementing the existing inventory of ETEC vaccine antigens. One potential drawback of O-PS as an ETEC vaccine target is the fact there are more than 78 O serogroups. Fortuitously, however, as few as 10 serogroups (O6, O8, O9, O25, O27, O78, O128, O148, O153 and O159) account for >75% of known ETEC isolates ^20^, suggesting that a 10-valent conjugate vaccine could afford broad protection. This number is much more feasible for multivalent ETEC vaccine development, especially considering that some of the most effective licensed conjugates achieve a valency of >10 by attachment of distinct polysaccharides from the most important serogroups (*e.g*., Prevnar13, Prevnar20). It is also worth noting that multivalency is not a unique challenge for an O-PS-based vaccine. There are 25 distinct CF/CS antigens identified to date that would need to be combined in some manner in order to achieve ~75% coverage of all isolates expressing the most common colonization factors ^10^. Hence, both O-PS and protein antigen-based vaccine candidates face complicated paths to a broadly protective, multivalent vaccine.

The ETEC O148 and O78 structures are now part of an ever-expanding list of polysaccharides that can be transferred to acceptor proteins by the *Cj*PglB biocatalyst. *Cj*PglB is well known for its remarkably relaxed oligosaccharide substrate specificity that allows transfer of diverse Und-PP-linked glycans including numerous different O-PS structures ^26, 45^. The ability of *Cj*PglB to site-specifically modify diverse acceptor proteins is aided by the introduction of a genetically encoded *N-*linked glycosylation tag that can be appended N- or C-terminally in single or multiple copies, or can be inserted at internal locations in the acceptor protein ^33^. Here, the introduction of four tandemly repeated DQNAT motifs at the C-termini of CRM_197_ and PD facilitated their use as acceptor protein substrates for *Cj*PglB and significantly expanded the set of carrier proteins available for bioconjugation, which historically has focused on a narrow set of carriers that are not currently used in any licensed vaccines – most notably *Pseudomonas aeruginosa* exotoxin A (ExoA) ^25^. Importantly, the in-built flexibility of bacterial OSTs like *Cj*PglB together with programmable glycosylation motifs makes it straightforward to produce an array of custom vaccines comprised of different polysaccharide/protein combinations.

As mentioned above, manufacturing of traditional conjugate vaccines is a complex, multistep process that is expensive, time consuming, and low yielding ^24^. The Centers for Disease Control and Prevention (CDC) cost per dose for conjugate vaccines ranges from ~$9.50 for the *H. influenzae* vaccine ActHIB up to ~$75 and ~$118 for the meningococcal vaccine Menactra and pneumococcal vaccine Prevnar 13, respectively ^46^. The cell-based and cell-free strategies described here represent greatly simplified alternatives for biomanufacturing conjugate vaccines whereby metabolically engineered *E. coli* or their cell-free extracts are exploited for one-step production of an unlimited and renewable supply of pure conjugate vaccine product. Because of the dramatic simplification of the process, bioconjugation strategies are anticipated to allow scalable production of large quantities of conjugates at a much more affordable cost, which is especially important for achieving sustained impact on global public health. For large-scale *E. coli*-based production, recent technoeconomic analysis indicates that the manufacturing costs (costs of goods sold, COGS) for a biologic antiviral, Griffithsin (GRFT), are as low as $3.43/g assuming production of >24,000 kg GRFT/yr ^47^. These authors concluded that, based on an estimated dose of 12–24 mg, it would be possible to provide ~1 billion doses/yr at a COGS of about $0.08/dose. While economics for a conjugate vaccine would be different due to the smaller dose and need for fewer total doses overall, we anticipate that a large-scale, *E. coli*-based manufacturing process for conjugates would offer a highly competitive cost per dose. Likewise, our own economic analysis of cell-free production revealed that iVAX reactions are inexpensive, costing ~$5/mL for raw materials. This cost can be decreased by approximately four-fold to ~$1.40/mL (~$0.50 per 24 μg dose) by optimizing the cell-free extract formulation to use the low-cost energy substrate maltodextrin in place of the significantly more expensive phosphorylated secondary energy substrate phosphoenolpyruvate (PEP) ^42^. Additional advantages of cell-free systems for conjugate production include that they can be: (i) distributed through freeze drying ^48^ followed by simple rehydration at the point of use ^27, 49^; (ii) linearly scaled from 1 nL to 100 L ^50^ for accelerated process development; and (iii) rapidly customized and reconfigured for product switching ^49, 51^. Taken together, these features have the potential to advance new paradigms in decentralized manufacturing of conjugate vaccines that we are continuing to explore for ETEC and other high-priority pathogens.

## MATERIALS AND METHODS

### Bacterial strains and plasmids

All strains used in this study are provided in **Supplementary Table 1**. Briefly, *E. coli* strain DH5α was used for all molecular biology including plasmid cloning and isolation. *E. coli* W3110 was used for expressing O-PS on lipid A-core and displaying O-PS molecules on the cell surface while *E. coli* CLM24 ^26^ was used as the host strain for expressing glycoengineered conjugates using intact cells. CLM24 is a derivative of W3110 that carries a deletion in the gene encoding the WaaL ligase, thus facilitating the accumulation of preassembled glycans on Und-PP as substrates for *Cj*PglB-mediated protein glycosylation. CLM24 Δ*lpxM* was used as the source strain for expressing glycoengineered conjugates in cell-free reactions. CLM24 Δ*lpxM* lacks the gene encoding the lipid A acyltransferase LpxM, a deletion that yields a pentaacylated lipid A structure with significantly reduced toxicity ^52^. The ETEC strains B7A (serotype O148:H28; CS6, LT, STa) ^30^ and H10407 (serotype O78:H11; CFA/I, LT, STa) ^43^ were used for SBA and challenge studies as well as a source of LPS. These strains were chosen because both were previously used in human challenge trials ^31, 43^.

Plasmids used in the study are listed in **Supplementary Table 1**. Briefly, plasmids pTrc99A-ssDsbA-PD^4xDQNAT^ and pTrc99A-ssDsbA-CRM_197_^4xDQNAT^ 27 were used to express the PD and CRM_197_ acceptor proteins in the *E. coli* periplasm. These plasmids were constructed by PCR amplification of the ssDsbA-PD^4xDQNAT^ and ssDsbA-CRM_197_^4xDQNAT^ sequences from plasmids pTrc99S-ssDsbA-PD^4xDQNAT^ and pTrc99S-ssDsbA-CRM_197_^4xDQNAT^, respectively ^27^, and followed by ligation of the PCR products in pTrc99A. Plasmid sequences were confirmed by Sanger sequencing at the Genomics Facility of the Cornell Biotechnology Resource Center (BRC). Plasmids pMAF10 ^26^ and pMAF10^D54N/E316Q 34^ were used for cell-based expression of wild-type *Cj*PglB and an inactive D54N/E316Q mutant of *Cj*PglB, respectively. The bacterial O-PS biosynthetic pathway plasmids were pMW07-O78 ^27, 29, 42^ for expressing the O-PS of ETEC strain H10407 and pMW07-O148 ^28, 29^ for expressing the O-PS of ETEC strain B7A. Plasmids used for cell-free expression of conjugate vaccines included pSF-*Cj*PglB-LpxE for co-expressing CjPglB along with *Francisella tularensis* phosphatase LpxE that promotes monophosphorylation of lipid A ^27^ and pJL1-PD^4xDQNAT^ for expressing PD^4xDQNAT 27^.

### Cell-based glycoconjugate expression and purification

For cell-based glycoconjugate expression, plasmids pTrc99A-ssDsbA-PD^4xDQNAT^ and pTrc99A-ssDsbA-CRM_197_^4xDQNAT^ encoding conjugate carrier proteins preceded by the DsbA signal peptide for translocation to the periplasm were used to transform CLM24 cells carrying a bacterial O-PS pathway encoded on plasmid pMW07-O148 or pMW07-O78 and *Cj*PglB encoded on plasmid pMAF10 or pMAF10^D54N/E316Q^. Transformed cells were grown in 10 mL LB medium (10 g/L yeast extract, 5 g/L tryptone, 5 g/L NaCl) overnight at 37 °C. The next day, cells were subcultured into 1 L of LB and allowed to grow at 37 °C until the optical density at 600 nm (OD_600_) reached 0.6-0.8. The culture was then supplemented with 0.2% arabinose to induce expression of *Cj*PglB and grown at 30 °C for 16 h, after which 0.5 mM isopropyl-β-D-thiogalactopyranoside (IPTG) was added to induce expression of the conjugate carrier protein for an additional 8 h at 30 °C. The cells were harvested, and cell pellets were resuspended in lysis buffer (20 mM Tris-HCl, 200 mM NaCl, 10 mM imidazole; pH 7.5) at 2-5 mL buffer per gram wet weight. Cells were lysed using a EmulsiFlex-C5 homogenizer (Avestin) then centrifuged at 13,000 × g for 30 min. The lysate was filtered through a 0.45-μm syringe filter and loaded onto a gravity flow column containing Ni-NTA resin (Thermo Fisher Scientific) that was washed with 5-10 column volumes of wash buffer (20 mM Tris-HCl, 200 mM NaCl, 20 mM imidazole; pH 7.5). The resin and cell lysate supernatant were allowed to equilibrate for 30 min at 4 °C. After flowthrough of the supernatant, the resin was washed with 5 column volumes of wash buffer. The protein was eluted with 3 column volumes of elution buffer (20 mM Tris-HCl, 200 mM NaCl, 300 mM imidazole; pH 7.5). Eluted protein was dialyzed into sterile PBS, concentrated, and quantified measured by Bradford assay.

### Periplasmic extract preparation

Periplasmic extracts containing the expressed glycoproteins were prepared by centrifuging the induced cultures at 13,000 × g and 4 °C for 2 min. The resulting pellets were resuspended in 0.4 M L-arginine (Sigma-Aldrich; 100 μL 0.4 M L-arginine per 100 mL culture) and incubated at 4 °C for 1 h with gentle shaking at 10-min intervals. The resuspended pellets were then centrifuged as above to obtain the final periplasmic extracts in the supernatant.

### Cell-free extract preparation

Cell-free extracts were prepared as previously described ^42, 53^. Specifically, CLM24 Δ*IpxM* cells were transformed with both pSF-*Cj*PglB-LpxE and pMW07-O78 plasmids for the strain used to generate PD-O78, and only the pSF-*Cj*PglB-LpxE plasmid for the strain used to generate the negative control aglycosylated PD. Cells were grown in a Sartorius Stedim BIOSTAT Cplus bioreactor at the 10-L scale in 2xYTP media supplemented with carbenicillin at 100 μg/mL and chloramphenicol at 34 μg/ml or only carbenicillin at 100 μg/mL in the negative control extract. Cells were inoculated at OD_600_ ≈ 0.08 and induced at OD_600_ ≈ 1 with 0.02% arabinose to induce expression of *Cj*PglB and O-PS enzymes and harvested at OD_600_ ≈ 3. All subsequent steps were performed on ice unless otherwise stated. Cells were harvested by centrifugation at 5,000 × g for 15 min and then washed 3 times with S30 buffer (10 mM Tris acetate pH 8.2, 14 mM magnesium acetate, and 60 mM potassium acetate). Following washing, cells were pelleted at 7,000 × g for 10 min, then flash frozen and stored at −80 °C. For lysis, CLM24 Δ*IpxM* cells were resuspended in 1 mL/g S30 buffer then homogenized using an EmulsiFlex-C3 high-pressure homogenizer (Avestin) with 1 pass at a pressure of ~21,000 psig. Following lysis, cells were centrifuged for 12,000 × g for 10 min. Supernatant was then collected and incubated at 37°C for 1 h in a runoff reaction. Cells were then centrifuged once more at 10,000 × g for 10 min and the supernatant was flash frozen and stored at –80 °C as the final extract.

### Cell-free protein synthesis reactions

For cell-free synthesis of PD-O78 and aglycosylated PD, reactions were prepared at the 5-mL scale in 50 mL conical tubes. Reactions producing PD-O78 used extract enriched with both *Cj*PglB and ETEC-O78 O-PS, while reactions productingg aglycosylated PD used extract enriched only with PglB. Each reaction was prepared as described previously ^54^ to contain 3.33 ng/μL pJL1-PD-4xDQNAT plasmid and 30% (vol./vol.%) extract in addition to: 10 mM magnesium glutamate (Sigma, 49605), 10 mM ammonium glutamate (Biosynth, FG28929), 130 mM potassium glutamate (Sigma, G1501), 1.2 mM adenosine triphosphate (Sigma A2383), 0.85 mM guanosine triphosphate (Sigma, G8877), 0.85 mM uridine triphosphate (Sigma U6625), 0.85 mM cytidine triphosphate (Sigma, C1506), 0.034 mg/mL folinic acid, 0.171 mg/mL *E. coli* tRNA (Roche 10108294001), 2 mM each of 20 amino acids, 30 mM phosphoenolpyruvate (PEP, Roche 10108294001), 0.4 mM nicotinamide adenine dinucleotide (Sigma N8535-15VL), 0.27 mM coenzyme-A (Sigma C3144), 4 mM oxalic acid (Sigma, PO963), 1 mM putrescine (Sigma, P5780), 1.5 mM spermidine (Sigma, S2626), 57 mM HEPES (Sigma, H3375), and 15-20 μg/mL T7. Reactions were then lyophilized for 16-20 hours using a VirTis Benchtop Pro Lyophilizer (SP scientific). Fully lyophilized reactions were rehydrated with 5 mL nuclease-free water and incubated at 30°C for one hour to synthesize the carrier protien (PD). After one hour of protein synthesis, glycosylation was initiated by supplementing 25 mM MnCl_2_ and 0.1 % wt/vol DDM. Reactions were incubated for one more hour at 30 °C, then were centrifuged at 16,000 × g for 15 minutes. The His-tagged carrier protein was then purified from the soluble cell-free reactions using Ni-NTA affinity resin as previously described ^42^.

### Western blot analysis

Cell-based glycoconjugate samples were run on NuPAGE 4-12% Bis-Tris gels (Invitrogen). Following electrophoretic separation, proteins were transferred from gels onto 0.45-μm Immobilon-P polyvinylidene difluoride membranes (PVDF) using a mini blot module (Thermo Fisher Scientific) according to the manufacturer’s instructions. Membranes were washed twice with TBS buffer (80 g/L NaCl, 20 g/L KCl, and 30 g/L Tris-base) followed by incubation for 1 h in blocking solution (50 g/L non-fat milk in TBS). After blocking, membranes were washed three times in TBS-T (TBS with 0.05% (v/v%) Tween-20) with a 5-min incubation between each wash. For fluorescence-based detection of immunoblots, membranes were probed with both an anti–6x-His tag antibody (R&D Systems, Cat # MAB050; diluted 1:7,500) and anti–ETEC O148 antibody (Abcam, Cat # ab78827; diluted 1:1,000) or anti–ETEC O78 antibody (Abcam, Cat # ab78826; diluted 1:1,000) in 1X TBS-T with 5% (w/v) BSA. Probing of membranes was performed overnight at 4 °C with gentle rocking, after which membranes were washed with TBS-T as described above and probed with fluorescently labeled secondary antibodies for 1 h at room temperature in 1X TBS-T with 5% (w/v) nonfat dry milk. The membrane was washed for 5 min with TBS-T and then imaged using a ChemiDoc XRS+System (Bio-Rad). For chemiluminescence-based detection of immunoblots, membranes were probed with an anti–6x-His tag antibody (Abcam, Cat # ab9108; diluted 1:7,500) and then with the corresponding anti-mouse HRP-conjugated secondary antibody (Abcam, Cat # ab205718; diluted 1:7,500). Another membrane was separately probed with anti-ETEC O148 antibody (Abcam, Cat # ab78827; diluted 1:1,000) or anti-ETEC O78 antibody (Abcam, Cat # ab78826; diluted 1:1,000) in 1X TBS-T with 5% (w/v) BSA and then with anti-rabbit HRP-conjugated secondary antibody (Abcam, Cat # ab205718; diluted 1:7,500). For signal visualization, membranes were briefly incubated at room temperature with Western ECL substrate (Bio-Rad) and imaged using a ChemiDoc XRS+System (Bio-Rad).

Cell-free samples were run on 4-12% Bis-Tris gels with SDS-MOPS running buffer supplemented with NuPAGE antioxidant. Samples were then transferred to PVDF 0.45-μm membranes (Millipore, USA) for 55 min at 80 mA per blot using a semi-dry transfer cell. Membranes were blocked for 1 h at room temperature or overnight at 4 °C in Intercept Blocking Buffer (Licor). Primary antibodies used were anti-His antibody (Abcam, Cat # ab1187; diluted 1:7,500) or anti-ETEC-O78 antibody (Abcam, Cat # ab78826; diluted 1:2,500) in Intercept blocking buffer with 0.2% (v/v) Tween 20, and membranes were incubated for 1 h at room temp or overnight at 4 °C. The secondary antibody used was a fluorescent goat anti-rabbit antibody (Licor, Cat # GAR-680RD; diluted 1:10,000) in Intercept blocking buffer, 0.2% (v/v) Tween 20 and 0.1% (w/v) SDS for both anti-His and anti-ETEC-O78 blots. Blots were washed 6 times for 5 min after each of the blocking, primary, and secondary antibody incubations using PBS-T. Blots were imaged with Licor Image studio.

### Dot blot analysis

To detect cell-surface expression of ETEC O148 O-PS, overnight cultures of the following strains were grown: *E. coli* strains W3110 and CLM24 without a plasmid, W3110 carrying empty pMW07, W3110 and CLM24 carrying pMW07-O148, and ETEC strain B7A. A total of 2 μl containing an equivalent amount of each strain, as well as LPS extracted from B7A cells, were spotted onto a nitrocellulose membrane. The membrane was allowed to dry, and then non-specific sites were blocked by soaking in 5% (w/v) BSA in TBS-T for 1 h at room temperature, followed by incubation for 30 min with anti–ETEC O148 antibody (Abcam, Cat # ab78827; diluted 1:1,000) in 0.1% (w/v) BSA in TBS-T. The membrane was washed three times with TBS-T, then incubated with secondary antibody conjugated to HRP. After three TBS-T washes, the membrane was incubated with Western ECL substrate (Bio-Rad) and imaged using a ChemiDocTM XRS+System (Bio-Rad). An identical protocol was followed for detection of cell-surface expression of ETEC O78 O-PS.

### Immunization

Groups of four 6-week-old female BALB/c mice (Harlan Sprague Dawley) were immunized s.c. with 50 μL of sterile PBS (pH 7.4, Fisher Scientific) or formulations containing either aglycosylated PD or PD-O148 conjugate. The amount of antigen in each preparation was normalized such that ~25-50 μg of these proteins was administered per injection. The purified protein groups were formulated in sterile PBS and mixed with an equal volume of Adju-Phos aluminium phosphate adjuvant (InvivoGen) before injection. Mice were boosted 21 and 42 days after the initial immunization. For antibody titering, blood was taken on days 0, 35, and 49 via submandibular collection, as well as at the study termination on day 56 via cardiac puncture. Sera were isolated from the collected blood draws after centrifugation at 5,000 × g for 10 min and stored at −20°C. For bacterial killing assays, final blood serum collections for all the mice within each group were pooled. The protocol number for the animal trial was 2012-0132 and was approved by the Institutional Animal Care and Use Committee (IACUC) at Cornell University.

### Serum antibody titering

Serum antibody titers were determined by ELISA using LPS derived from ETEC strains as immobilized antigen. Specifically, O148 and O78 LPS molecules were prepared from ETEC strains B7A and H10407, respectively, by hot phenol water extraction and DNase I (Sigma) and proteinase K (Invitrogen) treatment, as described elsewhere ^17^. Briefly, extracted LPS samples were purified using PD-10 desalting columns packed with Sephadex G-25 resin (Cytiva), and concentrations were determined using a purpald assay ^55^. 96-well plates (MaxiSorp; Nunc Nalgene) were incubated with 0.5 μg/mL of purified LPS diluted in PBS, pH 7.4, 25 μL/well, at 4 °C overnight. Plates were incubated with 50 μL blocking buffer (5% (w/v) nonfat dry milk (Carnation) in PBS) overnight at 4 °C, then washed three times with 200 μL PBS-T (PBS, 0.05% (v/v) Tween 20) per well. Serum samples isolated from the collected blood draws of immunized mice were appropriately serially diluted in triplicate in blocking buffer and added to the plates for 2 h at 37 °C. Plates were washed three times with PBS-T (+ 0.03% BSA (w/v)), then incubated for 1 h at 37 °C in the presence of a horseradish peroxidase– conjugated goat anti-mouse IgG antibody (Abcam, Cat # ab97265; diluted 1:25,000). After three PBS-T + 0.3% BSA washes, 50 μL of 3,3′-5,5′-tetramethylbenzidine substrate (1-Step Ultra TMB-ELISA; Thermo Fisher Scientific) was added to each well, and the plates were incubated at room temperature in the dark for 30 min. The reaction was stopped by adding 50 μL of 2 M H_2_SO_4_, and absorbance was measured at a wavelength of 450 nm using a FilterMax F5 microplate spectrophotometer (Agilent). Serum antibody titers were determined by measuring the lowest dilution that resulted in signals that were 3 standard deviations above the background controls of no serum. Statistical significance was determined by unpaired *t* test with Welch’s correction using GraphPad Prism 9 for MacOS (Version 9.2.0).

### SBA

A modified version of a previously described SBA method was followed ^56^. ETEC B7A cells were grown overnight from a frozen glycerol stock, then seeded 1:20 in LB medium. Log-phase grown bacteria were harvested, adjusted to an OD_600_ of 0.1, then further diluted 1:5,000 in Hanks’ Balanced Salt Solution with 0.5% (w/v) BSA (Sigma Aldrich). Assay mixtures were prepared in 96-well microtiter plates by combining 20 μL of serially diluted heat-inactivated test serum (dilutions ranging from 10^0^-10^4^), and 10 μL of diluted bacterial suspension. After incubation with shaking for 60 min at 37 °C, 10 μL of active or inactive complement source was added to each well, to a final volume percent of 25% (v/v). Heat-inactivated complement was prepared by thawing an aliquot of active pooled human complement serum (Innovative Research, ICSER1ML), incubating in a 56 °C water bath for 30 min, and cooling at room temperature. Assay plates were incubated with shaking at 37 °C for 60–90 min, then 10 μL was plated from each well (diluted to 50 μL in LB) on LB agar plates. Serum samples were tested and plated in duplicate, and colonies were counted (Promega Colony Counter) after 16–18 h of incubation at 30 °C. CFUs were counted for each individual serum dilution, and SBA titers were determined by calculating percent survival at various serum dilutions. Data was plotted as percentage survival versus serum dilution.

### ETEC infection

Groups of eight 6-week-old female BALB/c mice (Harlan Sprague Dawley) were immunized s.c. with 50 μL of sterile PBS (pH 7.4, Fisher Scientific) or formulations containing aglycosylated PD or PD-O148 conjugate, according to the 49-day immunization schedule described above. At 7 days post-vaccination (48 h prior to challenge infection), mice received streptomycin (5 g/L) and fructose (6.7% (w/v)) in the drinking water to eradicate normal flora and fecal pellets were collected. Food was withheld 12 h prior to challenge infection and replaced with sterile water without antibiotics. Famotidine (50 mg/kg) (Sigma-Aldrich) was then administered 2 h prior to challenge infection to neutralize gastric acid. At 9 days post-vaccination, mice were subsequently infected with a 200-μL inoculum containing ~1×10^4^ CFU of ETEC strain B7A, administered by gavage with a feeding needle directly introduced in the stomach via the esophagus. After challenge infection, mice were checked daily for 3 days, and fecal pellets were collected and stored at −20°C for further analysis. Mice were sacrificed 72 h post-infection. All procedures were carried out in accordance with protocol 2012-0132 approved by the Cornell University Institutional Animal Care and Use Committee.

### Quantitative real-time PCR analysis of ETEC burden

DNA from fecal pellets of individual mice was extracted from thawed stool samples using a QIAamp DNA stool kit (Qiagen) following the manufacturer’s instructions. To enhance extraction of pathogen DNA, stool samples were first vigorously homogenized with ~300 mg of 1.0-mm-diameter zirconia beads (Bio Spec) using a Mini-Bead Beater (BioSpec). After extraction, DNA was eluted in elution buffer and stored at −20°C. Stool DNA and tissue were analyzed for the ETEC B7A-specific heat-labile enterotoxin LT encoded by the *eltA* gene to determine the levels of shedding of the organism in stool. Quantification of ETEC was performed by qPCR using Taq DNA polymerase, as described elsewhere ^57, 58^, using the following conditions: preheating at 95 °C for 5 min, denaturation at 95 °C for 30 s, annealing at 58 °C for 30 s, elongation at 72 °C for 1 min. PCR was performed for 55-60 cycles for maximal saturation of signal with final extension at 72 °C for 7 min in a 7500 Fast Real-Time PCR System (Applied Biosystems). The primer sequences used were: *eltA* forward 5′-TTCCCACCGGATCACCAA −3′ and *eltA* reverse 5′-CAACCTTGTGGTGCATGATGA - 3′, along with a custom Taqman Probe (Thermo Fisher Scientific) 5’-CTTGGAGAGAAGAACCCT-3’ labeled with FAM (6-carboxyfluorescein) at the 5’ end and MGB at the 3′ end. Reactions with no DNA template were also included as controls. Reaction components were combined in MicroAmp 96-well reaction plates (Applied Biosystems) and plates were centrifuged at ~500 × g for 1 min before each reaction.

### Statistical analysis and reproducibility

To ensure robust reproducibility of all results, experiments were performed with at least three biological replicates and at least three technical measurements. Sample sizes were not predetermined based on statistical methods but were chosen according to the standards of the field (at least three independent biological replicates for each condition), which gave sufficient statistics for the effect sizes of interest. All data were reported as average values with error bars representing standard error of the mean (SEM). Statistical significance was determined by unpaired *t* test with Welch’s correction (\**p* < 0.05, \*\**p* < 0.01; ns, not significant). All graphs were generated using Prism 9 for MacOS version 9.2.0. No data were excluded from the analyses. The experiments were not randomized. The Investigators were not blinded to allocation during experiments and outcome assessment.

## Acknowledgements

This work was supported by the Bill and Melinda Gates Foundation (OPP1217652 to M.P.D. and M.C.J.), the Defense Threat Reduction Agency (grants HDTRA1-15-10052 and HDTRA1-20-10004 to M.P.D. and M.C.J.), and the National Science Foundation (grant CBET-1605242 to M.P.D. and grants CBET-1936823 and MCB-1413563 to M.P.D. and M.C.J.). A.J.W. was supported by a Cornell Presidential Postdoctoral Fellowship and K.F.W. was supported by the National Defense Science and Engineering (NDSEG) Fellowship Program (ND-CEN-013-096) sponsored by the Army Research Office.

## Author Contributions

Conceptualization: A.J.W., K.F.W., M.C.J., Y.-F.C. and M.P.D.

Methodology: A.J.W., P.D., K.F.W., J.L., J.-J.L., D.A.W., S.E.S. and Y.-F.C.

Investigation: A.J.W., P.D., K.F.W., J.L., J.-J.L., D.A.W., S.E.S. and Y.-F.C.

Visualization: M.P.D.

Supervision: M.C.J., Y.-F.C. and M.P.D. Writing - original draft: A.J.W. and M.P.D.

Writing - review & editing: A.J.W., P.D., K.F.W., M.C.J., Y.-F.C. and M.P.D.

## Data Availability

All data needed to evaluate the conclusions in the paper are present in the paper and/or the Supplementary Information.

## REFERENCES

1. Qadri, F., Svennerholm, A.M., Faruque, A.S. & Sack, R.B. Enterotoxigenic Escherichia coli in developing countries: epidemiology, microbiology, clinical features, treatment, and prevention. Clin Microbiol Rev 18, 465–483 (2005).

2. Khalil, I.A. et al. Morbidity and mortality due to shigella and enterotoxigenic Escherichia coli diarrhoea: the Global Burden of Disease Study 1990-2016. Lancet Infect Dis 18, 1229–1240 (2018).

3. Lamberti, L.M., Bourgeois, A.L., Fischer Walker, C.L., Black, R.E. & Sack, D. Estimating diarrheal illness and deaths attributable to Shigellae and enterotoxigenic Escherichia coli among older children, adolescents, and adults in South Asia and Africa. PLoS Negl Trop Dis 8, e2705 (2014).

4. Kotloff, K.L. et al. Burden and aetiology of diarrhoeal disease in infants and young children in developing countries (the Global Enteric Multicenter Study, GEMS): a prospective, case-control study. Lancet 382, 209–222 (2013).

5. Guerrant, R.L., DeBoer, M.D., Moore, S.R., Scharf, R.J. & Lima, A.A. The impoverished gut--a triple burden of diarrhoea, stunting and chronic disease. Nat Rev Gastroenterol Hepatol 10, 220–229 (2013).

6. Kotloff, K.L. et al. Global burden of diarrheal diseases among children in developing countries: Incidence, etiology, and insights from new molecular diagnostic techniques. Vaccine 35, 6783–6789 (2017).

7. WHO 2020 WHO Product Development for Vaccines Advisory Committee (PDVAC). Update on development of Enterotoxigenic E. coli (ETEC) vaccines. (2020).

8. Bourgeois, A.L., Wierzba, T.F. & Walker, R.I. Status of vaccine research and development for enterotoxigenic Escherichia coli. Vaccine 34, 2880–2886 (2016).

9. Riddle, M.S., Chen, W.H., Kirkwood, C.D. & MacLennan, C.A. Update on vaccines for enteric pathogens. Clin Microbiol Infect 24, 1039–1045 (2018).

10. Fleckenstein, J.M. Confronting challenges to enterotoxigenic Escherichia coli vaccine development. Front Trop Dis 2(2021).

11. Khalil, I. et al. Enterotoxigenic Escherichia coli (ETEC) vaccines: Priority activities to enable product development, licensure, and global access. Vaccine 39, 4266–4277 (2021).

12. Fleckenstein, J.M. et al. Molecular mechanisms of enterotoxigenic Escherichia coli infection. Microbes Infect 12, 89–98 (2010).

13. Darsley, M.J. et al. The oral, live attenuated enterotoxigenic Escherichia coli vaccine ACE527 reduces the incidence and severity of diarrhea in a human challenge model of diarrheal disease. Clin Vaccine Immunol 19, 1921–1931 (2012).

14. Harro, C. et al. Live attenuated enterotoxigenic Escherichia coli (ETEC) vaccine with dmLT adjuvant protects human volunteers against virulent experimental ETEC challenge. Vaccine 37, 1978–1986 (2019).

15. Behrens, R.H. et al. Efficacy and safety of a patch vaccine containing heat-labile toxin from Escherichia coli against travellers’ diarrhoea: a phase 3, randomised, double-blind, placebo-controlled field trial in travellers from Europe to Mexico and Guatemala. Lancet Infect Dis 14, 197–204 (2014).

16. Chakraborty, S. et al. Human Experimental Challenge With Enterotoxigenic Escherichia coli Elicits Immune Responses to Canonical and Novel Antigens Relevant to Vaccine Development. J Infect Dis 218, 1436–1446 (2018).

17. Svennerholm, A.M. et al. Induction of mucosal and systemic immune responses against the common O78 antigen of an oral inactivated ETEC vaccine in Bangladeshi children and infants. Vaccine 40, 380–389 (2022).

18. Fleckenstein, J., Sheikh, A. & Qadri, F. Novel antigens for enterotoxigenic Escherichia coli vaccines. Expert Rev Vaccines 13, 631–639 (2014).

19. Raetz, C.R. & Whitfield, C. Lipopolysaccharide endotoxins. Annu Rev Biochem 71, 635–700 (2002).

20. Wolf, M.K. Occurrence, distribution, and associations of O and H serogroups, colonization factor antigens, and toxins of enterotoxigenic Escherichia coli. Clin Microbiol Rev 10, 569–584 (1997).

21. Begum, Y.A. et al. Shift in phenotypic characteristics of enterotoxigenic Escherichia coli (ETEC) isolated from diarrheal patients in Bangladesh. PLoS Negl Trop Dis 8, e3031 (2014).

22. Avery, O.T. & Goebel, W.F. Chemo-Immunological Studies on Conjugated Carbohydrate-Proteins : Ii. Immunological Specificity of Synthetic Sugar-Protein Antigens. J Exp Med 50, 533–550 (1929).

23. Rappuoli, R. Glycoconjugate vaccines: Principles and mechanisms. Sci Transl Med 10(2018).

24. Frasch, C.E. Preparation of bacterial polysaccharide-protein conjugates: analytical and manufacturing challenges. Vaccine 27, 6468–6470 (2009).

25. Kay, E., Cuccui, J. & Wren, B.W. Recent advances in the production of recombinant glycoconjugate vaccines. NPJ Vaccines 4, 16 (2019).

26. Feldman, M.F. et al. Engineering N-linked protein glycosylation with diverse O antigen lipopolysaccharide structures in Escherichia coli. Proc Natl Acad Sci U S A 102, 3016–3021 (2005).

27. Stark, J.C. et al. On-demand biomanufacturing of protective conjugate vaccines. Sci Adv 7(2021).

28. Celik, E. et al. Glycoarrays with engineered phages displaying structurally diverse oligosaccharides enable high-throughput detection of glycan-protein interactions. Biotechnol J 10, 199–209 (2015).

29. Chen, L. et al. Outer membrane vesicles displaying engineered glycotopes elicit protective antibodies. Proc Natl Acad Sci U S A 113, E3609–3618 (2016).

30. DuPont, H.L. et al. Pathogenesis of Escherichia coli diarrhea. N Engl J Med 285, 1–9 (1971).

31. Levine, M.M. et al. Immunity to enterotoxigenic Escherichia coli. Infect Immun 23, 729–736 (1979).

32. Chen, M.M., Glover, K.J. & Imperiali, B. From peptide to protein: comparative analysis of the substrate specificity of N-linked glycosylation in C. jejuni. Biochemistry 46, 5579–5585 (2007).

33. Fisher, A.C. et al. Production of secretory and extracellular N-linked glycoproteins in Escherichia coli. Appl Environ Microbiol 77, 871–881 (2011).

34. Ollis, A.A., Zhang, S., Fisher, A.C. & DeLisa, M.P. Engineered oligosaccharyltransferases with greatly relaxed acceptor-site specificity. Nat Chem Biol 10, 816–822 (2014).

35. Borrow, R., Balmer, P. & Miller, E. Meningococcal surrogates of protection-- serum bactericidal antibody activity. Vaccine 23, 2222–2227 (2005).

36. Yang, X. et al. Serum antibodies protect against intraperitoneal challenge with enterotoxigenic Escherichia coli. J Biomed Biotechnol 2011, 632396 (2011).

37. Sears, K.T. et al. Bioactive Immune Components of Anti-Diarrheagenic Enterotoxigenic Escherichia coli Hyperimmune Bovine Colostrum Products. Clin Vaccine Immunol 24(2017).

38. Fleckenstein, J.M. & Rasko, D.A. Overcoming Enterotoxigenic Escherichia coli Pathogen Diversity: Translational Molecular Approaches to Inform Vaccine Design. Methods Mol Biol 1403, 363–383 (2016).

39. Bolick, D.T. et al. Critical Role of Zinc in a New Murine Model of Enterotoxigenic Escherichia coli Diarrhea. Infect Immun 86(2018).

40. Medeiros, P. et al. A bivalent vaccine confers immunogenicity and protection against Shigella flexneri and enterotoxigenic Escherichia coli infections in mice. NPJ Vaccines 5, 30 (2020).

41. Chakraborty, S. et al. Impact of lower challenge doses of enterotoxigenic Escherichia coli on clinical outcome, intestinal colonization and immune responses in adult volunteers. PLoS Negl Trop Dis 12, e0006442 (2018).

42. Warfel, K.F. et al. A low-cost, thermostable, cell-free protein synthesis platform for on demand production of conjugate vaccines. bioRxiv, 2022.2008.2010.503507 (2022).

43. Levine, M.M. et al. Lack of person-to-person transmission of enterotoxigenic Escherichia coli despite close contact. Am J Epidemiol 111, 347–355 (1980).

44. O’Neill, J. (2014).

45. Wacker, M. et al. Substrate specificity of bacterial oligosaccharyltransferase suggests a common transfer mechanism for the bacterial and eukaryotic systems. Proc Natl Acad Sci U S A 103, 7088–7093 (2006).

46. Prevention, C.f.D.C.a. (2019).

47. Decker, J.S., Menacho-Melgar, R. & Lynch, M.D. Low-Cost, Large-Scale Production of the Anti-viral Lectin Griffithsin. Front Bioeng Biotechnol 8, 1020 (2020).

48. Hunt, J.P., Yang, S.O., Wilding, K.M. & Bundy, B.C. The growing impact of lyophilized cell-free protein expression systems. Bioengineered 8, 325–330 (2017).

49. Pardee, K. et al. Portable, On-Demand Biomolecular Manufacturing. Cell 167, 248–259 e212 (2016).

50. Zawada, J.F. et al. Microscale to manufacturing scale-up of cell-free cytokine production--a new approach for shortening protein production development timelines. Biotechnol Bioeng 108, 1570–1578 (2011).

51. Adiga, R. et al. Point-of-care production of therapeutic proteins of good-manufacturing-practice quality. Nat Biomed Eng 2, 675–686 (2018).

52. Needham, B.D. et al. Modulating the innate immune response by combinatorial engineering of endotoxin. Proc Natl Acad Sci U S A 110, 1464–1469 (2013).

53. Jaroentomeechai, T. et al. Single-pot glycoprotein biosynthesis using a cell-free transcription-translation system enriched with glycosylation machinery. Nat Commun 9, 2686 (2018).

54. Hershewe, J.M. et al. Improving cell-free glycoprotein synthesis by characterizing and enriching native membrane vesicles. Nat Commun 12, 2363 (2021).

55. Lee, C.H. & Tsai, C.M. Quantification of bacterial lipopolysaccharides by the purpald assay: measuring formaldehyde generated from 2-keto-3-deoxyoctonate and heptose at the inner core by periodate oxidation. Anal Biochem 267, 161–168 (1999).

56. Valentine, J.L. et al. Immunization with Outer Membrane Vesicles Displaying Designer Glycotopes Yields Class-Switched, Glycan-Specific Antibodies. Cell Chem Biol 23, 655–665 (2016).

57. Liu, J. et al. A laboratory-developed TaqMan Array Card for simultaneous detection of 19 enteropathogens. J Clin Microbiol 51, 472–480 (2013).

58. Panchalingam, S. et al. Diagnostic microbiologic methods in the GEMS-1 case/control study. Clin Infect Dis 55 Suppl 4, S294–302 (2012).

