## Supplementary Table and Figures for "A low-cost recombinant glycoconjugate vaccine confers immunogenicity and protection against enterotoxigenic *Escherichia coli* infections in mice"

**Supplementary Table 1.** Strains and plasmids used in this study.

| **Strain or Plasmid** | **Description** | **Source or Reference** |
|---|---|---|
| *E. coli* DH5α | *supE44* Δ(*lacZYA-argF*)*U196* (Φ80Δ*lacZM15*) *hsdR17 recA1 endA1 gyrA96 thi-1 relA1* | Lab stock |
| *E. coli* W3110 | $F^-$ $\lambda^-$ *mcrA mcrB* IN(*rrnD-rrnE*)1 | Lab stock |
| *E. coli* CLM24 | *E. coli* W3110 with deletion in the *waaL* gene encoding the O-antigen ligase | 1 |
| *E. coli* CLM24 Δ*lpxM* | *E. coli* CLM24 with knockout of the *lpxM* gene encoding the lipid A acyltransferase | 2 |
| ETEC strain B7A | O148:H28; CS6, LT, STa | 3 |
| ETEC strain H10407 | O78:H11; CFA/I, LT, STa | 4 |
| pTrc99A-ssDsbA-PD$^{4xDQNAT}$ | *H. influenzae* protein D modified at N-terminus with signal peptide derived from *E. coli* DsbA and modified at C-terminus with 4x repeat of DQNAT glycosylation sequon and a 6x-His tag in the pTrc99A expression vector, Amp$^R$ | This study |
| pTrc99A-ssDsbA-CRM$_{197}^{4xDQNAT}$ | Non-toxic mutant of *Corynebacterium diphtheriae* diphtheria toxin modified at N-terminus with signal peptide derived from *E. coli* DsbA and modified at C-terminus with 4x repeat of DQNAT glycosylation sequon and a 6x-His tag in the pTrc99A expression vector, Amp$^R$ | This study |
| pMAF10 | HA-tagged *Cj*PglB cloned in pMLBAD, Tmp$^R$ | 1 |
| pMAF10$^{D54N/E316Q}$ | HA-tagged mutant *Cj*PglB with D54N/E316Q mutations cloned in pMLBAD, Tmp$^R$ | 5 |
| pMW07-O148 | *E. coli* O148 O-PS gene cluster in the pMW07 expression vector, Cm$^R$ | 6, 7 |
| pMW07-O78 | *E. coli* O78 O-PS gene cluster in the pMW07 expression vector, Cm$^R$ | 2, 7, 8 |
| pJL1-PD$^{4xDQNAT}$ | PD carrier protein modified at C-terminus with 4x repeat of DQNAT glycosylation sequon and a 6x-His tag in the pJL1 expression vector, Kan$^R$ | 2 |
| pSF-*Cj*PglB-LpxE | *Cj*PglB along with *Francisella tularensis* LpxE that promotes monophosphorylation of lipid A, Amp$^R$ | 27 |

### References


1. Feldman, M.F. et al. Engineering *N*-linked protein glycosylation with diverse O antigen lipopolysaccharide structures in *Escherichia coli*. *Proc Natl Acad Sci U S A* **102**, 3016-3021 (2005).
2. Stark, J.C. et al. On-demand biomanufacturing of protective conjugate vaccines. *Sci Adv* **7** (2021).
3. DuPont, H.L. et al. Pathogenesis of *Escherichia coli* diarrhea. *N Engl J Med* **285**, 1-9 (1971).
4. Levine, M.M. et al. Lack of person-to-person transmission of enterotoxigenic *Escherichia coli* despite close contact. *Am J Epidemiol* **111**, 347-355 (1980).
5. Ollis, A.A., Zhang, S., Fisher, A.C. & DeLisa, M.P. Engineered oligosaccharyltransferases with greatly relaxed acceptor-site specificity. *Nat Chem Biol* **10**, 816-822 (2014).
6. Celik, E. et al. Glycoarrays with engineered phages displaying structurally diverse oligosaccharides enable high-throughput detection of glycan-protein interactions. *Biotechnol J* **10**, 199-209 (2015).
7. Chen, L. et al. Outer membrane vesicles displaying engineered glycotopes elicit protective antibodies. *Proc Natl Acad Sci U S A* **113**, E3609-3618 (2016).
8. Warfel, K.F. et al. A low-cost, thermostable, cell-free protein synthesis platform for on demand production of conjugate vaccines. *bioRxiv*, 2022.2008.2010.503507 (2022).

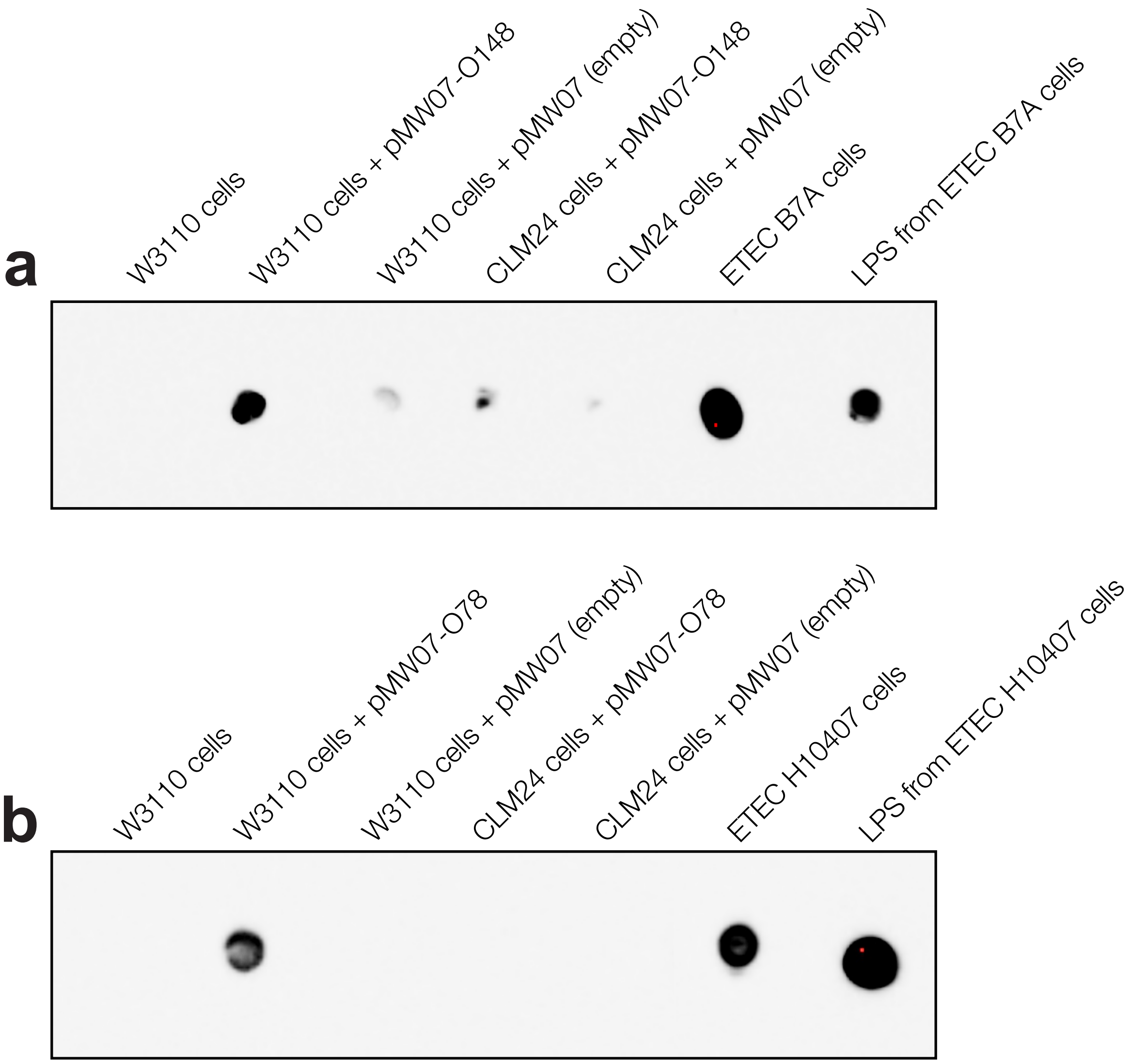


**Supplementary Figure 1. Biosynthesis of ETEC O-PS antigens in non-pathogenic *E. coli*.** Dot blot analysis of a whole cells and LPS derived from the strains indicated. A total of 2 µL of $OD_{600}$-matched cultures or LPS extracts were spotted on nitrocellulose membranes. Membranes were probed with (a) anti-ETEC O148 antibody or (b) anti-ETEC O78 antibody. Results are representative of three biological replicates.

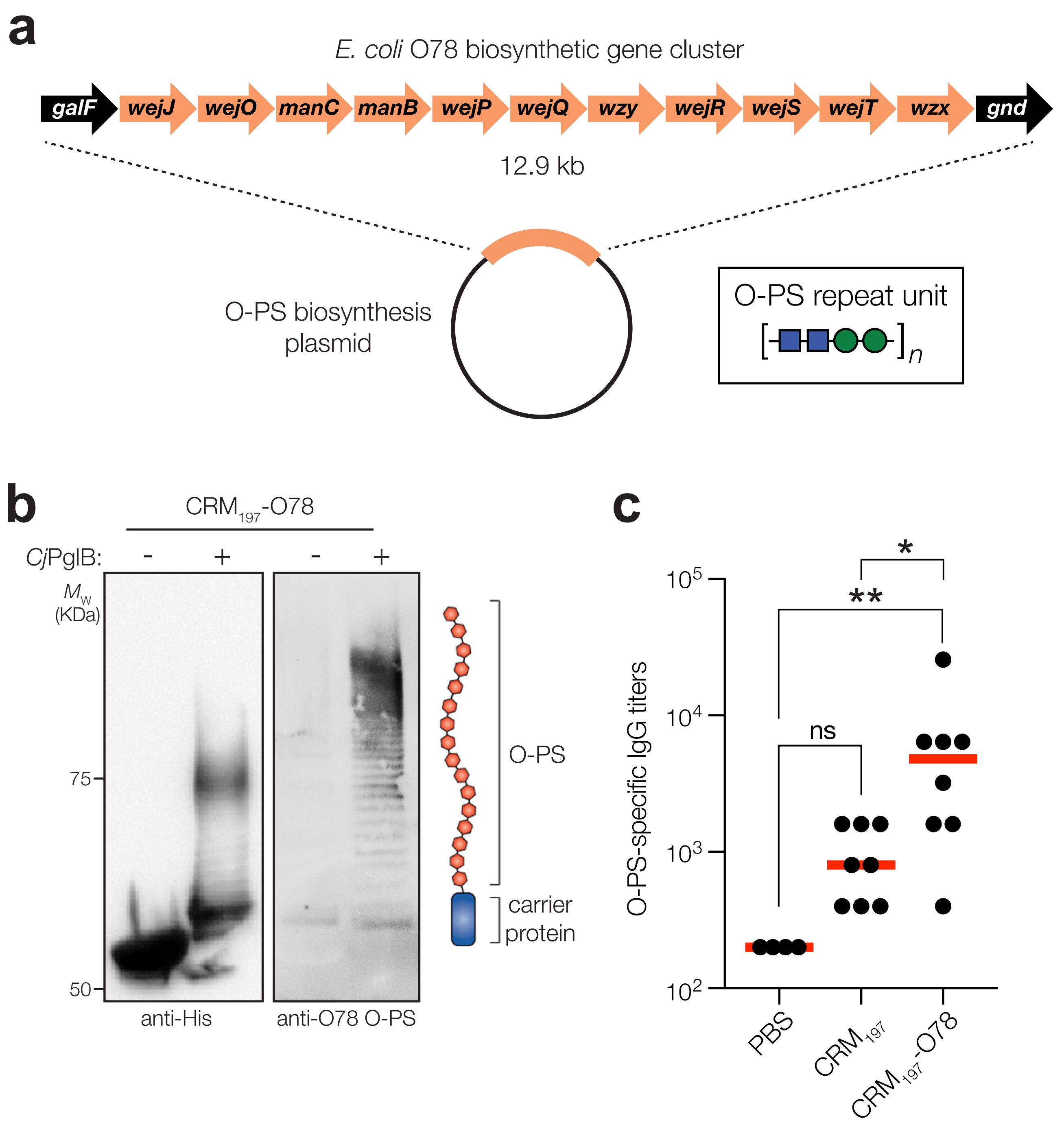


**Supplementary Figure 2. Biosynthesis of ETEC O148 O-PS antigen in non-pathogenic *E. coli*.** (a) Biosynthesis of ETEC O78 O-PS from plasmid pMW07-O78, which encodes the entire O-PS locus from ETEC strain H10407 (serotype O78:H11) between *galF* and *gnd*. (b) Immunoblot analysis of purified carrier proteins derived from *E. coli* CLM24 Δ*lpxM* cells carrying a plasmid encoding $CRM_{197}^{4xDQNAT}$ along with plasmid pMW07-O78 encoding the ETEC O78 O-PS biosynthetic pathway and plasmid pMAF10 encoding wild-type *Cj*PglB (wt) or an inactive mutant of *Cj*PglB (mut) as indicated. Blots were probed with anti-His antibody to detect acceptor proteins and anti-ETEC O78 antibody to detect O-PS. Images depict aglycosylated and multiply glycosylated forms of $CRM_{197}^{4xDQNAT}$. Molecular weight ($M_W$) markers are indicated on the left. Results are representative of three biological replicates. (c) O-PS-specific IgG titers in day 56 serum of individual mice (black circles) and mean titers of each group (red lines) as determined by ELISA with LPS derived from ETEC H10407 as immobilized antigen. Groups of three BALB/c mice were immunized s.c. with 100 µL PBS alone or PBS containing 50 µg of either aglycosylated $CRM_{197}^{4xDQNAT}$ carrier protein or glycosylated $CRM_{197}^{4xDQNAT}$ bearing ETEC O78 O-PS adjuvanted with aluminium phosphate adjuvant. Mice were boosted on days 21 and 42 with identical doses of each immunogen. Statistical significance was determined by unpaired *t* test with Welch's correction (*$p < 0.05$; **$p < 0.01$; ns, not significant).
